# Merging OpenLifeData with SADI services using Galaxy and Docker

**DOI:** 10.1101/013615

**Authors:** Mikel Egañna Aranguren

## Abstract

Semantic Web technologies have been widely applied in Life Sciences, for example by data providers like OpenLifeData and Web Services frameworks like SADI. The recent OpenLifeData2SADI project offers access to the OpenLifeData data store through SADI services. This paper shows how to merge data from OpenLifeData with other extant SADI services in the Galaxy bioinformatics analysis platform, making semantic data amenable to complex analyses, as a worked example demonstrates. The whole setting is reproducible through a Docker image that includes a pre-configured Galaxy server with data and workflows that constitute the worked example.

## INTRODUCTION

The idea behind a 3rd generation Semantic Web is to codify information on the Web in a way that machines can directly access and process it, opening possibilities for automated analysis. Even though the Semantic Web has not yet been fully implemented, it has been used in the life sciences, in which Semantic Web technologies are used to integrate data from different resources with disparate schemas [Good and Wilkinson (2006)]. The Semantic Web is based on a set of standards proposed by the W3C (WWW Consortium^1^), including the following elements:

### RDF (Resource Description Framework)

RDF^2^ is a machine-readable data representation language based on the triple, *i.e*., data is represented in a subject-predicate-object structure (*e.g*. “Cyclin participates in Cell cycle”, Figure 1), in which the predicate and object (“participates in Cell cycle”) describe a property of the subject (“Cyclin”). A collection of connected triples forms a graph. In a graph, there are entities that are the object in one triple and the subject in another, connecting triples. Graphs are stored in data bases called triple stores.

**Figure 1.**
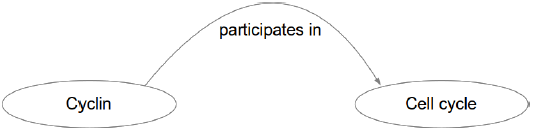
RDF triple. The predicate (“participates in”) goes from subject (“Cyclin”) to object (“Cell cycle”).

### SPARQL (SPARQL Protocol and RDF Query Language)

SPARQL^3^ is a query language to extract data from RDF graphs.

### OWL (Web Ontology Language)

OWL^4^ is a knowledge representation language for asserting general ideas about data (*e.g*. “A protein participates in at least one biological process”), using complex axioms so that automated reasoning can be applied. Therefore, OWL is used to create ontologies that codify the consensus of a community about their knowledge area. In an OWL ontology, entities are connected through logical axioms, and there are different types of entities: individuals are the actual instances of data (*e.g*. “Cyclin”, “Mikel”, *etc*.); properties link individuals (*e.g*. “Mikel lives in Bilbo”); classes are sets of individuals that fulfill certain conditions (*e.g*. “Protein”, “Human”, *etc*.).

The backbone of the Semantic Web stack is the fact that URIs^5^ (Uniform Resource Identifiers) are used to identify entities (OWL classes, instances and properties; RDF subjects, predicates and objects). This allows one to refer to entities from external resources over the Web, *e.g*. in an RDF triple, the subject might be indicated by a URI from one resource and the predicate and object by a URI from a different resource.

The most widely used implementation of SemanticWeb technologies in the life sciences and elsewhere is the set of principles from Linked Data for publishing data on the Web^6^:

1. Identify every data item (entity or relation between entities) with a URI.
2. Make those URIs HTTP resolvable (dereferenceable).
3. Provide information in an open standard when an entity is requested by HTTP, preferably in RDF. The format to provide should be defined by content negotiation between the client and the server (*e.g*. RDF for an automatic agent, HTML for a human user), so that the entity and its representations are decoupled in different URIs.
4. Assure that the information provided contains typed relations to other entities, so that the agent can browse through those relations and discover new information, analogous to regular Web browsers.

Linked Data has already shown its value as a means for data publication in a machine-readable and Web resolvable fashion, opening new and interesting possibilities [Aranguren et al. (2014a)]. Hence, important data providers in the life sciences have implemented Linked Data solutions, such as UniProt^7^, EBI RDF^8^, and OpenLifeData^9^, contributing to the growth of the Linked Open Data (LOD) cloud^10^.

On the other hand, in addition to data storage, Semantic Web standards have also been used for creating Semantic Web Services through the Semantic Automated Discovery and Integration (SADI) design patterns [Wilkinson et al. (2011)]. Recently, the OpenLifeData2SADI project has used SADI to wrap the OpenLifeData data store, providing access to the whole data store through several thousand services [González et al. (2014)]. This paper shows how to combine OpenLifeData2SADI services with existing SADI services, using off-the-self tools from the popular Galaxy bioinformatics platform [Goecks et al. (2010)]. Additionally, a worked example is provided as a ready to use Docker image containing a pre-configured SADI-Galaxy server, data and an appropriate workflow, making the procedure trivially reproducible. This approach provides multiple advantages and its reproducibility allows the potential to examine a wide variety of modifications.

## TECHNICAL BACKGROUND

### SADI services

SADI is a Semantic Web standards-based set of design patterns for providing web services. SADI does not require any new technology nor scheme for fully functional deployment (not even for the messaging infrastructure). SADI uses off-the-self, already tested technologies (URI, RDF, and OWL) to provide all the necessary functionality. In a SADI service, the data the server can consume is defined in an OWL class: the client uses automated reasoning to infer whether the input RDF is a member of the OWL class, and if so, it pushes the RDF to the service. Once the service has processed the input, the output is formed by the initial RDF instance augmented with additional triples. Effectively, SADI services produce chains of new Linked Data [González et al. (2014)].

### OpenLifeData2SADI

The Bio2RDF project captures existing data from different providers and publishes it again with normalised URIs and Linked Data support [Belleau et al. (2008)]. The OpenLifeData project wraps Bio2RDF and enhances its content negotiation functionality and axiomatic richness. Additionally, OpenLifeData2SADI offers access to OpenLifeData through different SADI services [González et al. (2014)]. That means that the actual data can be retrieved, in RDF, by simply calling the appropriate SADI service, and more importantly, the output RDF can be integrated with other Linked Data/SADI output.

### Galaxy

Galaxy is a Web server that offers an infrastructure in which biologists can analyse data through a consistent Web interface (Figure 2). A history of the performed tasks is stored so that workflows with common steps can be extracted from the history and re-run independently. The most common bioinformatics tools are already included in the Galaxy distribution, whereas new tools can be created by simply wrapping command line executables in Galaxy-compliant XML files. There are many public Galaxy servers^11^ and Galaxy can also be installed privately.

**Figure 2.**
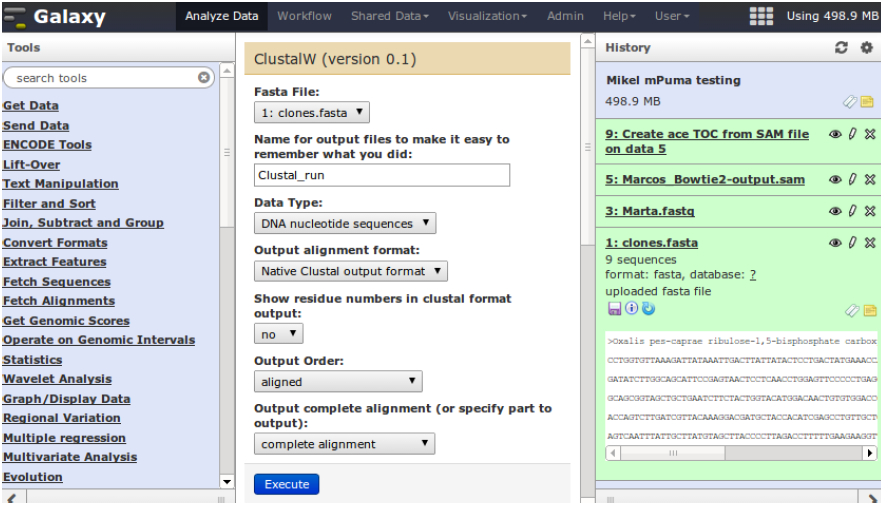
Galaxy main interface (Figure reproduced from [Aranguren et al. (2014b)]). Galaxy is a Web server with different interfaces: “Analyze data”, “Workflow”, “Shared data”, *etc*. The main interface, “Analyze data” (Shown in the Figure), is where data is analysed with different tools (left column), and a history is recorded (right column), so that workflows can be extracted (they will appear in the “Workflow” interface). In “Shared data” histories, data, and workflows can be shared between users and/or published.

### SADI-Galaxy

SADI-Galaxy^12^ is a Galaxy tool generator for SADI services [Aranguren et al. (2014b)]. It includes a SADI generic client that can invoke SADI services, given the service URL and an input RDF. It also includes an advanced functionality in that SADI-Galaxy can query a SADI registry^13^ and generate a Galaxy tool for each of the retrieved services, so that the user need not know the service URL in order to invoke it. The service simply appears as another Galaxy tool in the left column.

### Docker

Docker^14^ is a virtualisation engine and runtime. The key difference with Virtual Machines (VMs) is that a Docker image does not include the whole Operating System (OS), making images lighter (in the case of the host being a GNU/Linux system). Containers can be run, with the Docker engine, from pre-defined images. Docker is also a GitHub-like repository of images, so a developer can build an image with the desired computational environment (OS, libraries, configuration), software and data, starting from an already existing image (*e.g*. Ubuntu 14.04) and ship it to the repository. Then anyone can pull the image and run it, including the shipped software, without configuration/installation.

## WORKED EXAMPLE

### Merging OpenLifeData2SADI and SADI services in a single workflow

An example workflow shows how OpenLifeData2SADI and SADI services can be merged^15^ (Figures 3 and 4). The workflow answers the following question: Given a set of UniProt proteins, which ones are related to PubMed abstracts containing the term “brain”, and what are the known SNPs from dbSNP related to those proteins? The workflow starts from a simple list of UniProt IDs, and retrieves different datasets from a regular SADI service (to obtain SNPs) and a set of 3 OpenLifeData2SADI services (to obtain PubMed abstracts). The results are then merged and queried to obtain the SNPs of proteins that are related to PubMed abstracts that contain the term. The workflow involves six steps.

**Figure 3.**
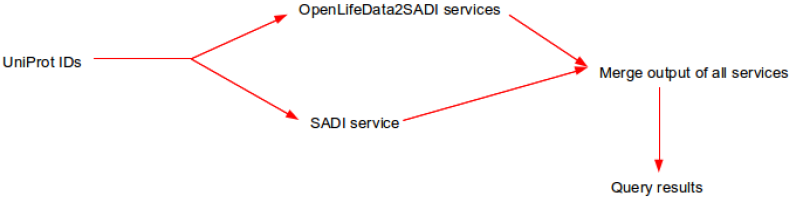
Conceptual representation of example workflow. The workflow starts from a set of UniProt IDs, and obtains information from OpenLifeData SADI services and regular SADI services. The output is merged in a single dataset and queried.

**Figure 4.**
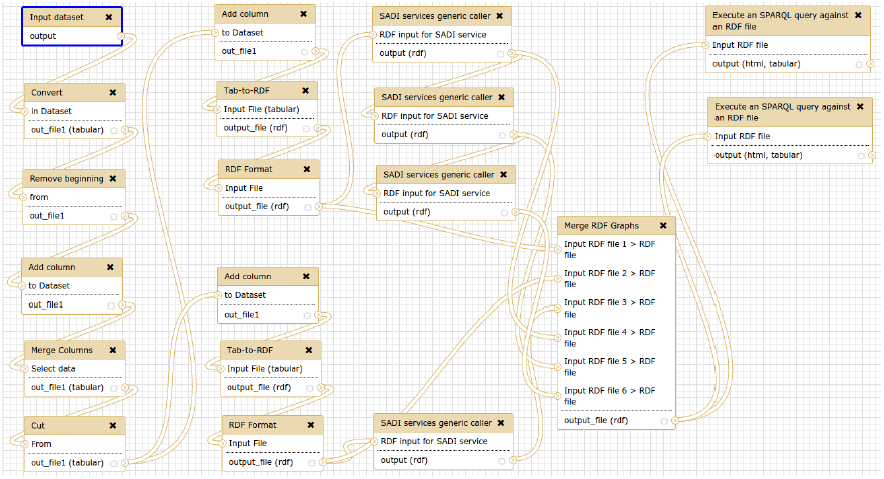
Screenshot of the actual Galaxy workflow that implements the general idea described in Figure 3. The workflow executes two groups of SADI services, and therefore the input UniProt IDs must be converted into two RDF datasets, but the first steps of the process are shared (from “Input Dataset” to “Cut”). Then the appropriate RDF triple is added to each UniProt ID (from “Add column” to “RDF Format”) and SADI services are called (“SADI services generic caller”). The output of the SADI services and the input RDF is merged in a single graph and then queried, producing the results in TSV format and HTML format.

#### 0.- Configure Galaxy tools

This step is not really necessary in this case, since the Docker image with the configured Galaxy instance is provided, but it is included for the sake of completeness. The SADI services for this workflow are executed with the generic client (OpenLifeData2SADI services) and with specific Galaxy tools generated from the SADI registry (The SADI service for getting the SNPs). In the latter case, the service tool was generated by retrieving the services compliant with the OWL class `UniProt record`^16^, retrieved with the following SPARQL query (See [Aranguren et al. (2014b)] for details on how to generate Galaxy tools with SADI-Galaxy):

~~~
PREFIX dc:
    <http://protege.stanford.edu/plugins/owl/dc/protege-dc.owl#>
PREFIX sadi: <http://sadiframework.org/ontologies/sadi.owl#>
PREFIX rdf: <http://www.w3.org/1999/02/22-rdf-syntax-ns#>
PREFIX serv: <http://www.mygrid.org.uk/mygrid-moby-service#>

SELECT ?s

WHERE {
 ?s rdf:type serv:serviceDescription .
 ?s rdf:type sadi:Service .
 ?s serv:hasOperation ?op .
 ?op serv:inputParameter ?input .
 ?input serv:objectType
 <http://purl.oclc.org/SADI/LSRN/UniProt_Record>
}
~~~

#### 1.- Obtain a list of UniProt IDs of interest

This can be done, for example, by simply uploading the list from the computer or importing it directly to Galaxy from Biomart:

~~~
Accession
Q03164
Q9UKA4
Q8TDM6
Q9NQT8
Q12830
Q9HCM3
Q8TF72
Q5H8C1
Q9UGU0
B2RWN9
A4UGR9
…
~~~

#### 2.- Convert the input to RDF

In order for the data to be consumed by the SADI services, it needs to be converted to RDF. Additionally, an `rdf:type` triple must be added to each ID asserting the OWL input class for each set of services, producing two different inputs from the same UniProt IDs list: rdf:type http://openlifedata.org/uniprotvocabulary:Resource for OpenLife-Data2SADI services and rdf:type http://purl.oclc.org/SADI/LSRN/UniProt Record for the service to retrieve SNPs (getdbSNPRecordByUniprotID). For example:

~~~
<?xml version="1.0" encoding="utf-8"?>
 <rdf:RDF xmlns:rdf="http://www.w3.org/1999/02/22-rdf-syntax-ns#">
  <rdf:Description rdf:about="http://openlifedata.org/uniprot:Q03164">
    <rdf:type rdf:resource="http://openlifedata.org/
      uniprot_vocabulary:Resource"/>
  </rdf:Description>
   <rdf:Description rdf:about="http://openlifedata.org/uniprot:Q9UKA4">
    <rdf:type rdf:resource="http://openlifedata.org/
      uniprot_vocabulary:Resource"/>
 </rdf:Description>
…
~~~

#### 3.- Send the appropriate input to services

Each of the RDF inputs is send to OpenLifeData2SADI services (3 services in a row) and getdbSNPRecordByUniprotID.

#### 4.- Merge the outputs and the inputs in a single RDF graph

Since SADI services keep track of the inputs by way of the URIs (The predicates are simply added to the input URIs and the input URIs are maintained in the output), the outputs of the services are merged with the inputs in a single graph.

#### 5.- Query the merged graph with SPARQL

In this case, the UniProt entries from the input set that are mentioned in a PubMed abstract with the term “brain” and their respective SNPs from dbSNP are retrieved with the following query (Figure 5):

~~~
PREFIX rdf: <http://www.w3.org/1999/02/22-rdf-syntax-ns#>
PREFIX rdfs: <http://www.w3.org/2000/01/rdf-schema#>
PREFIX sio_resource: <http://semanticscience.org/resource/>
SELECT ?protein ?label ?SNP
WHERE {
    ?protein rdf:type
       <http://openlifedata.org/uniprot_vocabulary:Resource> .
    ?protein sio_resource:SIO_000272 ?SNP .
    ?protein ?prot2hgnc ?hgnc .
    ?hgnc ?hgnc2omim ?omim .
    ?omim ?omim2pubmed ?pubmed .
    ?pubmed rdfs:label ?label .
    FILTER (regex (?label, ′brain′))
}
~~~

**Figure 5.**
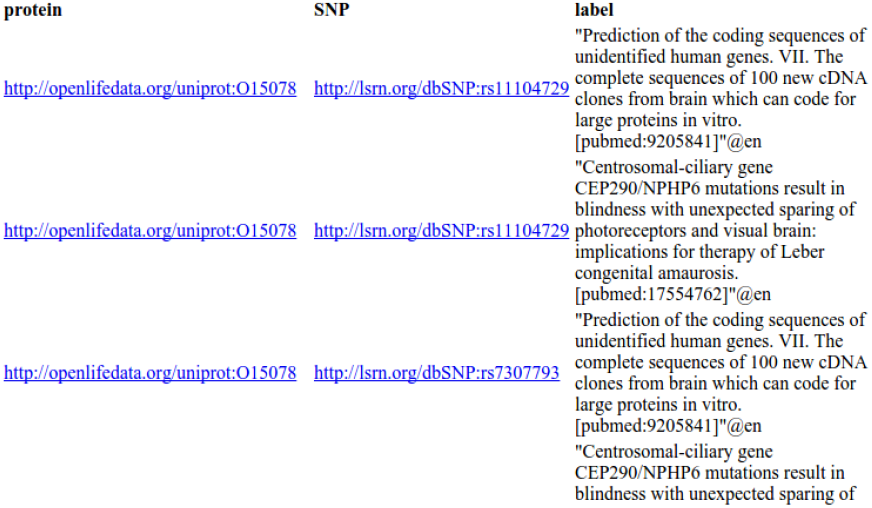
The result of the workflow is a list of PubMed abstracts with the term “Brain”, with related proteins and SNPs. The result can be displayed as HTML, for browsing the actual resources in their Webs, or TSV (Tab Separated Values), for downstream analysis in Galaxy.

### Reproducing the workflow through Galaxy and Docker

The workflow can be reproduced^17^ through the following Docker image^18^:

~~~
FROM ubuntu:14.04
MAINTAINER Mikel Egaña Aranguren <aray@kgi.edu>
RUN apt-get update && apt-get install -y git wget vim python
python-setuptools raptor2-utils libraptor2-0
RUN easy_install rdflib
RUN wget http://www.duinsoft.nl/pkg/pool/all/update-sun-jre.bin
RUN sh update-sun-jre.bin
RUN git clone
http://github.com/mikel-egana-aranguren/SADI-Galaxy-Docker.git
CMD /SADI-Galaxy-Docker/galaxy-dist/./run.sh
~~~

The image can be obtained from the Docker central registry^19^ by pulling it (Assuming a GNU/Linux system with the Docker engine installed):

~~~
$ docker pull mikeleganaaranguren/sadi-galaxy
~~~

The container is run by:

~~~
$ docker run -d -p 8080:8080 mikeleganaaranguren/sadi-galaxy
~~~

The command will start a container with Ubuntu 14.04, install and configure Galaxy^20^ and the appropriate libraries and tools, and will run Galaxy as a daemon on the port 8080^21^ (Figure 6). When the user logs in (a fake user has been defined, user, password: “useruser”), a history will appear, with the actual data and workflow (Figure 7). The workflow can be run again in order to reproduce the result (Figure 8).

**Figure 6.**
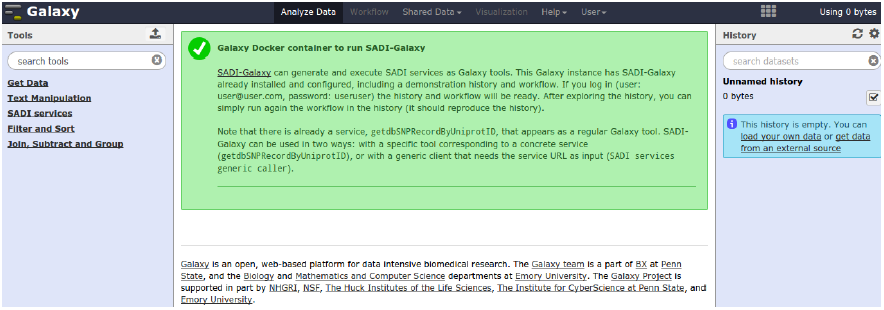
Galaxy server running as part of the Docker container. This Galaxy instance contains only the necessary tools to reproduce the use case (left column).

**Figure 7.**
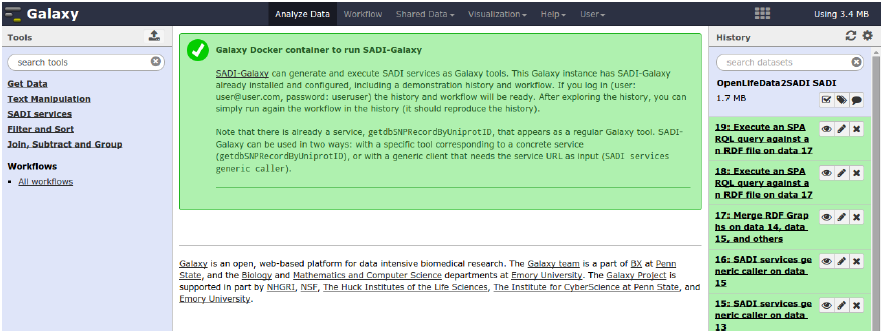
After logging in, a history with the use case appears (right column).

**Figure 8.**
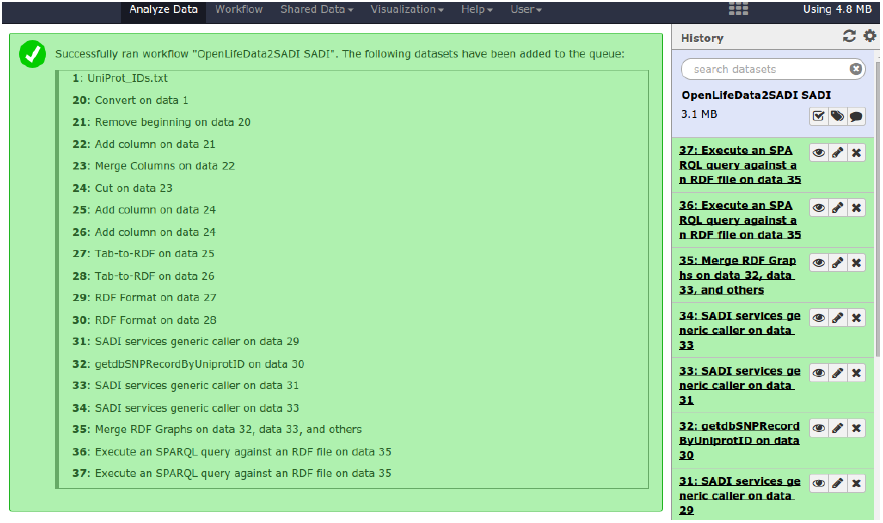
The workflow can be executed in the same history, using the first dataset as input. The workflow can be modified in any way, once it is run in the current history.

## DISCUSSION

### Data integration and manipulation through RDF and SADI

The data model behind RDF, the triple, is very simple, yet many different types of data can be modelled with it. Additionally, RDF uses URIs to identify the modelled entities. Therefore, the procedure described herein, *i.e*. accessing data from different resources and integrating it with SADI, is straight-forward and easily extendable, since it is based on moving RDF from one tool to the next. We can import only the RDF triples we need for our analysis from OpenLifeData, and integrate them with other datasets.

Also, the RDF data model makes input preparation for SADI a naive process, since TSV data can be easily mapped to RDF, and then mapped again to TSV for further processing, with standard Galaxy tools for text manipulation.

Finally, many URIs from the datasets described in this process are also URLs, *i.e*. they not only identify, they also locate entities on the Web. That means that, for example, many results of the final dataset are clickable (*e.g*. *e.g*. OpenLifeData URLs, see Figure 5).

### Reproducibility with Galaxy and Docker

Computational reproducibility is becoming an important area in life sciences [Garijo et al. (2013); Sandve et al. (2013)]. This use case demonstrates a procedure by which SADI services and OpenLifeData services can be merged in a completely reproducible fashion, by implementing reproducibility in two levels:

1. **- Virtualisation of computational environment (Operating System) through Docker**. Docker allows encapsulation of a complex environment with all the necessary data and software [Boettiger (2014)]. In this case, a Ubuntu 14.04 image is shipped, with SADI (and needed dependencies) and Galaxy completely configured, which means that the user need only log in into the Galaxy instance. The user does not have to deal with installing and configuring Galaxy and SADI-Galaxy.
2. **- Reproducibility of performed analyses through Galaxy**. Galaxy is a suitable environment for executing SADI services in a reproducible manner, since it provides an infrastructure in which workflow management, history and provenance, and data storage are pre-established^22^. That means that any SADI-Galaxy based analysis, if performed in a Galaxy instance, is easily reproducible. For example, the same workflow can be repeated every time OpenLifeData is updated and the workflow can be modified and/or fused with other workflows (Even if the workflows are not based on RDF/SADI).

## CONCLUSION

With this setup (SADI-Galaxy on top of Docker to call OpenLifeData2SADI and SADI services) one can simplify the calling of services and the integration of information in a completely reproducible manner. This facilitates the distribution of example workflows, which then can be modified in new contexts, providing a tool for distribution of case implementations in multi-platform environments. The use of the Galaxy interface additionally provides a single foundation for integration of services, the construction of RDF graphs and their subsequent querying. Here, the worked example provides a tangible illustration of the use of Semantic Web constructs and standards for the extraction of new information from disparate and independent services.

## ACKNOWLEDGEMENTS

Peter B. Pearman from the University of Basque Country corrected the writing and provided valuable comments about the content.

## Footnotes

1 http://www.w3.org/

2 www.w3.org/standards/techs/rdf

3 http://www.w3.org/standards/techs/sparql

4 http://www.w3.org/standards/techs/owl

5 http://tools.ietf.org/html/rfc3986

6 Adapted from http://www.w3.org/DesignIssues/LinkedData.html and [González et al. (2014)].

7 http://www.uniprot.org/

8 http://www.ebi.ac.uk/rdf/

9 http://openlifedata.org/

10 http://lod-cloud.net/

11 https://wiki.galaxyproject.org/PublicGalaxyServers

12 http://github.com/mikel-egana-aranguren/SADI-Galaxy

13 http://sadiframework.org/registry

14 http://www.docker.com/

15 This workflow, albeit new, builds upon workflows already presented in [Aranguren et al. (2014b)] and [González et al. (2014)].

16 http://purl.oclc.org/SADI/LSRN/UniProt_Record

17 In fact a user can simply create a new image pasting this instructions in a Docker file, since the appropriate Galaxy instance, configured and including data, is downloaded when cloning the GitHub repository (`RUN git clone` …).

18 http://github.com/mikel-egana-aranguren/SADI-Galaxy-Docker

19 http://registry.hub.docker.com/u/mikeleganaaranguren/sadi-galaxy/

20 The Galaxy instance is very simple, with only SADI-Galaxy, other necessary tools, and no PostgreSQL data base, since it is meant only as a demonstration. It should not be used for production and/or with big datasets.

21 The port 8080 of the host is forwarded to the port 8080 in the Docker container.

22 http://www.gigasciencejournal.com/series/Galaxy

